# Identification of a molecular fingerprint for synaptic glia

**DOI:** 10.1101/2020.03.13.991323

**Authors:** Ryan Castro, Thomas Taetzsch, Sydney K. Vaughan, Kerilyn Godbe, John Chappell, Robert Settlage, Gregorio Valdez

**Affiliations:** Department of Molecular Biology, Cellular Biology, and Biochemistry, Brown University, Providence, Rhode Island, USA; Fralin Biomedical Research Institute at Virginia Tech Carilion, Roanoke, Virginia, USA

## Abstract

The inability to specifically identify and manipulate synaptic glial cells remains a major obstacle to understanding fundamental aspects of synapse formation, stability and repair. Using a combinatorial gene expression approach, we discovered molecular markers that allow us to specifically label perisynaptic Schwann cells (PSCs), glial cells at neuromuscular synapses. Using these markers, we demonstrate that PSCs fully-differentiate postnatally and have a unique molecular signature that includes genes predicted and known to play critical roles at synapses. These findings will serve as a springboard for unprecedented approaches for studying molecular determinants of PSC differentiation and function at neuromuscular synapses and possibly synapse-associated glia throughout the CNS.

## Introduction

Synapses are formed, maintained and repaired through the coordinated actions of three distinct cellular components: the presynaptic and postsynaptic neuronal components, and the synaptic glia. A wealth of knowledge has accrued about the presynaptic and postsynaptic regions^1^ because they can be readily identified morphologically, and molecularly targeted at all stages of life and in a wide variety of conditions. In stark contrast, the identity and spatial distribution of synaptic glia are generally poorly defined and thus very little is known about the cellular and molecular mechanisms that govern their formation, differentiation and function at the synapse^2,3^. The slow progress in answering fundamental questions about synaptic glia can be primarily attributed to the lack of molecular tools with which to study them independently of other glial cells.

Although there exist a number of molecular markers that recognize subsets of glial cells throughout the nervous system^4^, none of these single markers show specificity for synaptic glia. We explored the possibility that synaptic glia may be distinguished by unique combinations of glial cell markers, determined by a cell-specific pattern of gene expression. Synaptic glia of both the central (CNS) and peripheral (PNS) nervous systems are generally thought to provide structural, functional and trophic support to the synapse. However, only the perisynaptic or terminal Schwann cells (PSCs)^3^ of the PNS, the non-myelinating Schwann cells of the neuromuscular junction (NMJ), can be readily identified by their unique morphology and presence at the synapse^3^. Despite this, the inability to selectively visualize and target PSCs remains a major obstacle to understanding the cellular and molecular rules that govern their differentiation and function at NMJs during development, following injury, in old age, and in diseases, such as ALS.

## Results and Discussion

We used a combinatorial gene expression approach to uncover markers specific for PSCs. We found that PSCs can be identified by a combination of two different glial marker proteins, calcium-binding protein B (S100B) and neuro-glia antigen-2 (NG2). S100B is present in all Schwann cells, while NG2, has been shown to be present in subsets of Schwann cells in addition to astrocytes, oligodendrocytes, pericytes, and endothelial cells^5^. To facilitate visualization of PSCs, we created a transgenic mouse line (referred herein as S100B-GFP;NG2-dsRed; Fig. 1A) by crossing transgenic lines in which either the NG2 promoter drives expression of dsRed ^6^ or the S100B promoter drives expression of GFP^7^. As expected, in the resulting S100B-GFP;NG2-dsRed double transgenic mouse line, dsRed labeled all NG2 positive cells (referred herein as NG2-dsRed+) and GFP labeled all Schwann cells (referred herein as S100B-GFP+) (Fig. 1B-C) in skeletal muscles. However, we found a select subset of glia specifically located at the NMJ positive for both S100B-GFP+ and NG2-dsRed+ (yellow cells in Fig. 1D). Based on the location and morphology of the cell body and its elaborations, we concluded that PSCs are the only cells expressing both S100B-GFP+ and NG2-dsRed+ in skeletal muscles. The co-expression of S100B-GFP+ and NG2-dsRed+ in PSCs had no apparent deleterious effect on either PSCs or the NMJ (Fig. 1E-F). Thus, we have discovered a unique combination of markers with which to readily identify and study the synaptic glia of the NMJ in a manner previously not possible.

**Fig. 1:**
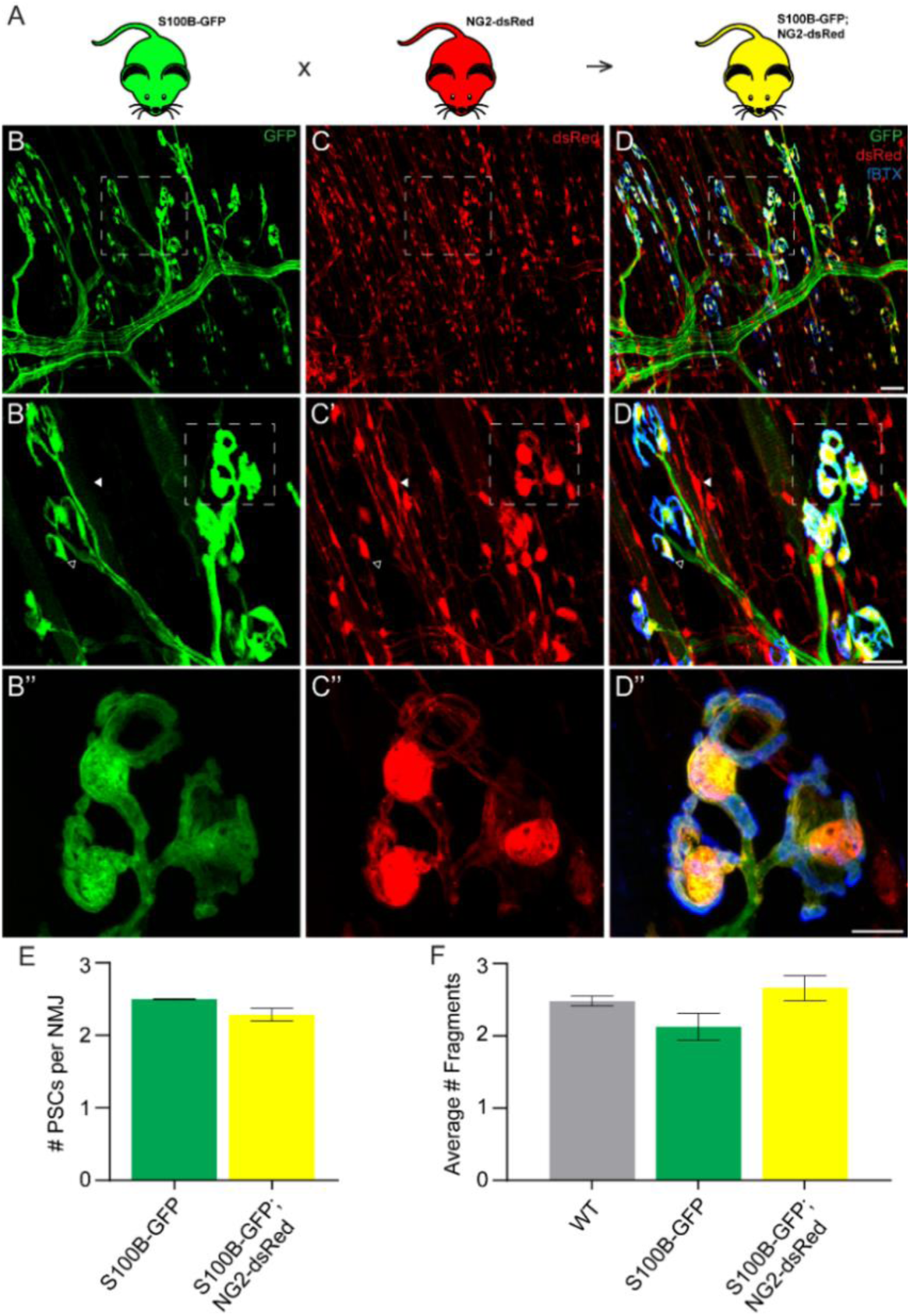
Co-expression of S100B and NG2 is unique to PSCs in muscles. In order to selectively label PSCs, S100B-GFP and NG2-dsRed transgenic mice were crossed to create S100B-GFP;NG2-dsRed mice (A). In the S100B-GFP mouse line, all Schwann cells express GFP (B, B’). In the NG2-dsRed mouse line, all NG2 positive cells express dsRed (C, C’). In S100B-GFP;NG2-dsRed mice, PSCs identified based on their unique morphology and location at NMJs, visualized here using fBTX to detect nAChRs (blue), are the only cells expressing both GFP and dsRed (D, D’). At non-synaptic sites, GFP-positive cells do not express dsRed (hollow arrow; B’, C’, D’) and dsRed-positive cells do not express GFP (filled arrow; B’, C’, D’). The co-expression of GFP and dsRed has no discernible negative effects on NMJ fragmentation or PSC number in the EDL muscle of young adult mice (E-F). The average number of PSCs per NMJ is unchanged between S100B-GFP mice and S100B-GFP;NG2-dsRed mice (E). The average number of nAChR clusters per NMJ is unchanged between wild-type, S100B-GFP, and S100B-GFP;NG2-dsRed animals (F). Error bar = standard error. Scale bar = 50 μm (D), 25 μm (D’), and 10 μm (D’’).

To begin to determine at which time PSCs acquire specific characteristics during development, we determined the earliest time point at which both S100B-GFP and NG2-dsRed were co-expressed in PSCs. We examined NMJs in the extensor digitorum longus (EDL) muscle of S100B-GFP;NG2-dsRed mice at various embryonic (E) and postnatal (P) stages (Fig. 2 and Fig. s1). This analysis revealed that NMJs associate exclusively with S100B-GFP+ cells at least until E18 (Fig. 2A-C). PSCs expressing both S100B-GFP+ and NG2-dsRed+ appear at the NMJ around P0 and become the only cell-type present at NMJs by P21 (Fig. 2A, C). In stark contrast, we failed to find cells expressing only NG2-dsRed+ at embryonic and postnatal NMJs. Thus, it appears that PSCs are defined by at least one key PSC-specific characteristic, NG2. To confirm that dsRed expression from the NG2 promoter denotes the temporal and spatial transcriptional control of the NG2 gene, we found NG2 protein present at postnatal but not embryonic NMJs (Fig. s2). The induced expression of NG2 in NMJ Schwann cells supports previous studies indicating that PSCs originate from Schwann cells^8^. The delayed expression of NG2 (Fig. 2A, C and Fig. s2) further indicates that fully-differentiated PSCs only become associated with NMJs after their initial formation.

**Fig. 2:**
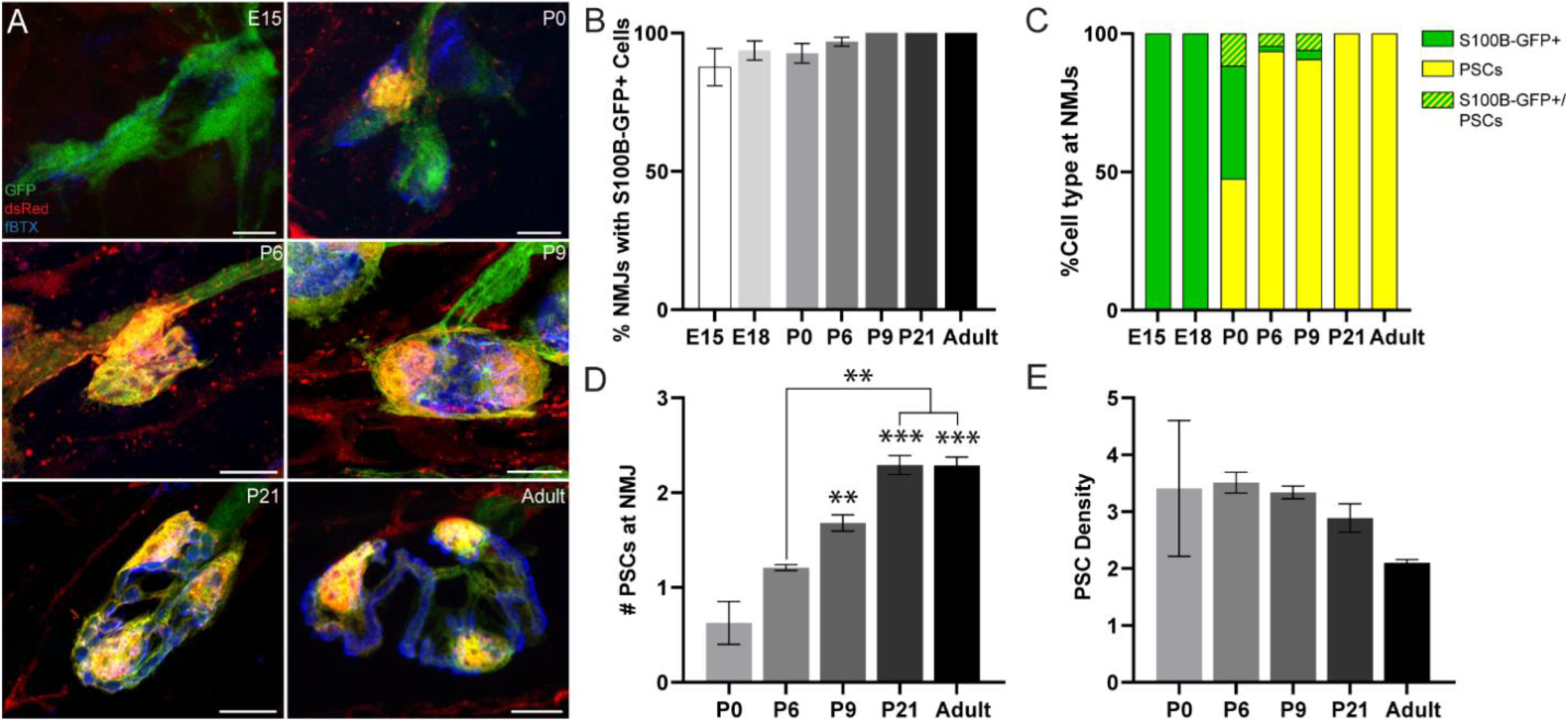
Analysis of PSCs at different developmental stages. NMJs are found associated exclusively with S100B-GFP+ cells between E15 and E18 (A-C). PSCs expressing both S100B-GFP+ and NG2-dsRed+ appear at the NMJ around P0 and become the only cell-type present at NMJs by P21 (A, C). The average number of PSCs per NMJ increases during development (D). When standardizing for the change in NMJ size during development, there is no difference in the density of PSCs at NMJs, represented as the number of PSCs per 500um^2^ of NMJ area (E). Error bar = standard error. Scale bar = 10 μm. ** = P<0.01; *** = P<0.001.

Previous studies relied solely on a combination of anatomical location and Schwann cell markers to make inferences about the number and spatial arrangement of PSCs at NMJs^9,10^. Such studies may have missed important relationships between PSCs and the NMJ particularly early in development when PSC appearance could not be easily discerned^11^. We thus reexamined the number of PSCs at developing and adult NMJs in the EDL muscle of S100B-GFP;NG2-dsRed mice. We found that the number of PSCs rapidly increased from P0 to P9 (Fig. 2A, D), a timespan when the NMJ undergoes rapid cellular, molecular and functional changes^12^. Highlighting the importance of specifically visualizing PSCs, we found NMJs populated by a combination of PSCs and S100B-GFP+ cells between P0 and P9 (Fig. 2C), further supporting the idea that PSCs are derived from Schwann cells at the NMJ. The number of PSCs reached an average of 2.3 per NMJ by P21 that remained unchanged in healthy young adult mice (Fig. 2A, D). A closer examination, however, revealed that while the number of PSCs varies across NMJs of different sizes, and in different muscle types (Fig. s3 A-D), their density remains unchanged (Fig. 2E & s3 E). These data demonstrate that the number of PSCs directly correlates with the size and not functional characteristics of individual NMJs.

The ability to distinguish PSCs from all other Schwann cells made it possible to identify genes either preferentially or specifically expressed in PSCs. For this, we deployed fluorescence-activated cell sorting (FACS) to separately isolate PSCs, single-labeled S100B-GFP+ Schwann cells, and single-labeled NG2-dsRed+ cells from juvenile S100B-GFP;NG2-dsRed transgenic mice (Fig. 3A, s4 A). Light microscopy and expression analysis of GFP and dsRed using quantitative PCR (qPCR) showed that only cells of interest were sorted (Fig. s4 B). We utilized RNA-sequencing (RNA-seq) to compare the transcriptional profile of PSCs to the other two cell types (Fig. 3A). This analysis revealed a unique transcriptional profile for PSCs (Fig. 3B). Using Ingenuity Pathway Analysis (IPA) we identified a number of genes that encode secreted and transmembrane proteins (Fig. 3C), raising the possibility that PSCs utilize them to promote the pruning, stability, repair and function of the NMJ. Furthermore, IPA identified synaptogenesis, glutamate receptor, and axon guidance signaling as top canonical pathways under transcriptional regulation (Fig. 3D). Altogether, we have shown that a unique gene expression signature distinguishes PSCs from all other Schwann cells. The manner in which PSCs use such gene products to shape the maturation and function of the neuromuscular synapse remains to be determined.

**Fig. 3:**
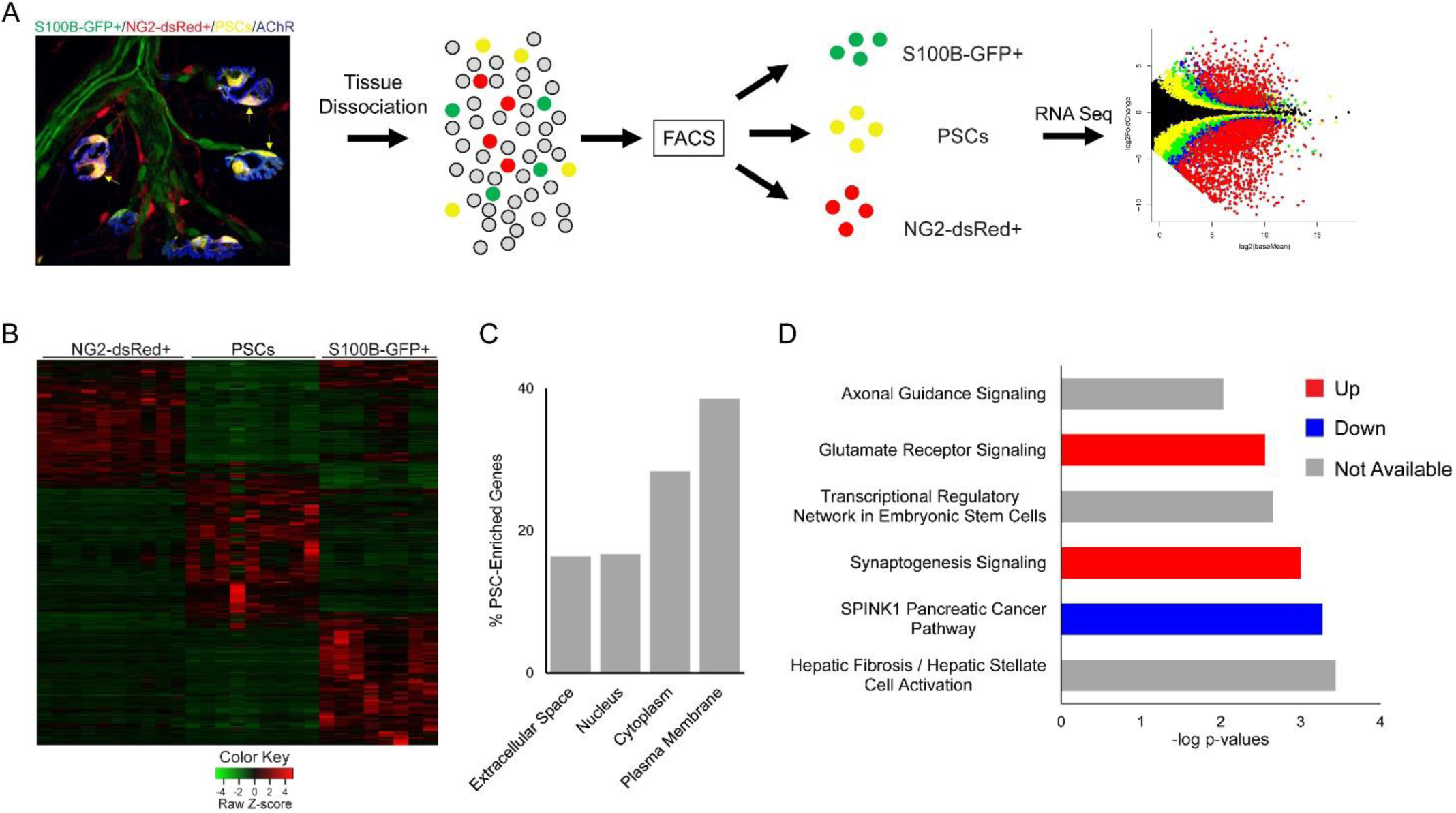
PSCs, S100B-GFP+ cells, and NG2-dsRed+ cells were analyzed by RNA-sequencing following FACS isolation from dissociated muscle tissue of juvenile S100B-GFP;NG2-dsRed mice (A). A heat map of RNA-seq results depicting genes with at least 5 counts and expression differences with a p-value of less than 0.01 between any 2 cell types reveals a distinct transcriptome in PSCs versus S100B-GFP+ and NG2-dsRed+ cells (B). Cellular location of genes enriched in PSCs versus S100B-GFP+, and NG2-dsRed+ cells according to Ingenuity Pathway Analysis. PSC enriched genes are defined by a minimum count of 5 in PSCs, as well as a 4-fold increase and a p-value of less than 0.05 versus other cell types (C). Synaptogenesis and axon guidance signaling are among the most influential signaling pathways in PSCs according to Ingenuity Pathway Analysis of genes enriched in PSCs versus S100B-GFP+, and NG2-dsRed+ cells (D). Error bar = standard error.

In summary, we have discovered molecular markers that allow us to specifically visualize, isolate, interrogate the transcriptome, and potentially alter the molecular composition of PSCs. With these tools, it is now possible to determine which cellular and molecular determinants are critical for PSC differentiation, maturation, and function at the NMJ. It will also allow us to ascertain the contribution of PSCs to NMJ repair following injury, degeneration during normal aging and the progression of neuromuscular diseases, such as Amyotrophic Lateral Sclerosis (ALS). Our strategy of specifically labeling synaptic glia, using combinations of protein markers uniquely expressed in this cell type, may serve as a springboard for unprecedented approaches for studying not only PSC function at the NMJ, but also synapse-associated glia throughout the CNS. Indeed, we have observed subsets of astrocytes in the brain that co-express both S100B and NG2, as has been previously reported in the context of a lineage tracing analysis^13^. Future studies will determine the generality of our approach in discerning the functional roles of synaptic glia in the development, maintenance and function of select synapses.

## Acknowledgments

We thank members of the Valdez laboratory for providing helpful comments throughout the course of this project.

## Materials and Methods

### Mice

S100B-GFP mice (B6;D2-Tg(S100B-EGFP)1Wjt/J) and NG2-dsRed mice (Tg(Cspg4-DsRed.T1)1Akik/J) were obtained from Jackson Labs (Bar Harbor, ME) and crossed to generate S100B-GFP;NG2-dsRed mice. Offspring were genotyped using Zeiss LSM900 to check for fluorescent labels. Postnatal mice older than 9 days of age were anesthetized and immediately perfused with 4% paraformaldehyde (PFA) overnight. Pups were anesthetized by isoflurane and euthanized by cervical dislocation prior to muscle dissociation. Adult mice were anesthetized using CO_2_ and then perfused transcardially with 10 ml of 0.1 M PBS, followed by 25 ml of ice-cold 4% PFA in 0.1 M PBS (pH 7.4). All experiments were carried out under NIH guidelines and animal protocols approved by the Brown University and Virginia Tech Institutional Animal Care and Use Committee.

### Immunohistochemistry and NMJ visualization

For NG2 immunohistochemistry (IHC), muscles were incubated in blocking buffer (5% lamb serum, 3% BSA, 0.5% Triton X-100 in PBS) at room temperature for 2h, incubated with anti-NG2 antibody (courtesy of Dr. Dwight Bergles) diluted at 1:250 in blocking buffer overnight at 4°C, washed 3 times with 0.1M PBS for 5 minutes. Muscles were then incubated with 1:1000 Alexa Fluor-488 conjugated anti-guinea pig antibody (A-11008, Invitrogen, Carlsbad, CA) and 1:1000 Alexa Fluor-555 conjugated α-bungarotoxin (fBTX; Invitrogen, B35451) in blocking buffer for 2h at room temperature and washed 3 times with 0.1M PBS for 5 minutes. For all other NMJ visualization, muscles were incubated in Alexa Fluor-647 conjugated α-bungarotoxin (fBTX; Invitrogen, B35450) at 1:1000 and 4’,6-Diamidino-2-Phenylindole, Dihydrochloride (DAPI; D1306, ThermoFisher, Waltham, MA) at 1:1000 in 0.1M PBS at 4°C overnight. Muscles were then washed with 0.1M PBS 3 times for 5 minutes each. Muscles were whole mounted using Vectashield (H-1000, Vector Labs, Burlingame, CA) and 24×50-1.5 cover glass (ThermoFisher).

### Confocal microscopy of PSCs and NMJs

All images were taken with a Zeiss LSM700, Zeiss LSM 710, or Zeiss LSM 900 confocal light microscope (Carl Zeiss, Jena, Germany) with a 20× air objective (0.8 numerical aperture), 40× oil immersion objective (1.3 numerical aperture), or 63× oil immersion objective (1.4 numerical aperture) using the Zeiss Zen Black software. Optical slices within the z-stack were taken at 1.00 μm or 2.00 μm intervals. High resolution images were acquired using the Zeiss LSM 900 with Airyscan under the 63× oil immersion objective in super resolution mode. Optical slices within the z-stack were 0.13 μm with a frame size of 2210 x 2210 pixels. Images were collapsed into a two-dimensional maximum intensity projection for analysis.

### Image analysis

#### NMJ size

To quantify the area of NMJs, the area of the region occupied by nAChRs, labeled by fBTX, was measured using ImageJ software. At least 100 nAChRs were analyzed for number of fragments, individual nicotinic acetylcholine receptor (nAChR) clusters, from each muscle to represent an individual mouse. At least 3 animals per age group were analyzed to generate the represented data.

#### Cells associated with NMJs

Cell bodies were visualized via GFP and/or dsRed signal, and were confirmed as cell bodies by the presence of a DAPI+ nucleus. The area of each cell body was measured by tracing the outline of the entire cell body using the freehand tool in ImageJ. To quantify the number of cells associated with NMJs, the number of cell bodies directly adjacent to each NMJ was counted. Every cell that overlapped with or directly abutted the fBTX signal was considered adjacent to the NMJ. At least 3 animals per age group were analyzed to generate the represented data. Cells were examined in at least 100 NMJs from each muscle to represent an individual mouse.

#### Spacing of PSCs at NMJs

NMJs were identified via fBTX signal. PSCs were identified by the colocalization of GFP, dsRed, and DAPI signal in addition to their location at NMJs. The area of each PSC and NMJ was measured. The linear distance from the center of each PSC soma to the center of the nearest PSC soma at a single NMJ was measured. The distances were then separated into 5μm bins and plotted in a histogram. All linear measurements were made using the line tool in the ImageJ software. At least 100 NMJs were analyzed from each muscle to represent an individual mouse.

### Muscle Dissociation and Fluorescence Activated Cell Sorting

Diaphragm, pectoralis, forelimb and hindlimb muscles were collected from p15-p21 S100B-GFP;NG2-dsRed mice. After removal of connective tissue and fat, muscles were cut into 5 mm^2^ pieces with forceps and digested in 2 mg/mL collagenase II (Worthington Chemicals, Lakewood, NJ) for 1h at 37°C. Muscles were further dissociated by mechanical trituration in Dulbecco’s modified eagle medium (Life Technologies, Carlsbad, CA) containing 10% horse serum (Life Techonoligies) and passed through a 40 um filter to generate a single cell suspension. Excess debris was removed from the suspension by centrifugation in 4% BSA followed by a second centrifugation in 40% Optiprep solution (Sigma-Aldrich, St. Louis, MO) from which the interphase was collected. Cells were diluted in FACS buffer containing 1mM EDTA, 25mM Hepes, 1% heat inactivated fetal bovine serum (Life Technologies), in Ca^2+^/Mg^2+^ free 1X Dulbecco’s phosphate buffered saline (Life Technologies). FACS was performed with a Sony SH800 Cell Sorter (Sony Biotechnology, San Jose, CA). Sorting gates were set at the lowest fluorescence threshold at which the sorted cell population was 100% pure and confirmed with dsRed and GFP qPCR (Fig. 3-s1B).

### RNA-seq and qPCR

RNA was isolated from S100B-GFP^+^, NG2-dsRed^+^, or S100B-GFP^+^/NG2-dsRed^+^ cells following FACS with the PicoPure RNA Isolation Kit (ThermoFisher). The maximum number of cells that could be collected by FACS following dissociation of muscles collected from one mouse was used as a single replicate. On average, a single replicate consisted of 7,500 cells. RNA seq was performed by Genewiz on 12 replicates per cell type. Following sequencing, data were trimmed for both adaptor and quality using a combination of ea-utils and Btrim^14,15^. Sequencing reads were aligned to the genome using Tophat2/HiSat2^16^ and counted via HTSeq ^17^. QC summary statistics were examined to identify any problematic samples (e.g. total read counts, quality and base composition profiles (+/- trimming), raw fastq formatted data files, aligned files (bam and text file containing sample alignment statistics), and count files (HTSeq text files). Following successful alignment, mRNA differential expression were determined using contrasts of and tested for significance using the Benjamini-Hochberg corrected Wald Test in the R-package DESeq2^18^. Failed samples were identified by visual inspection of pairs plots and removed from further analysis resulting in the following number of replicates for each cell type: NG2-dsRed^+^, 10; S100B-GFP^+^, 7; NG2-dsRed^+^/S100B-GFP^+^, 9. Functional and pathway analysis was performed using Ingenuity Pathway Analysis (QIAGEN Inc., https://www.qiagenbio-informatics.com/products/ingenuity-pathway-analysis). Quantitative reverse transcriptase PCR (qPCR) was performed on 6 replicates of each cell type. Reverse transcription was performed with iScript (Bio-Rad, Hercules, CA) and was followed by a preamplification PCR step with SsoAdvanced PreAmp Supermix (Bio-Rad) pSrior to qPCR using iTAQ SYBR Green and a CFX Connect Real Time PCR System (Bio-Rad). Relative expression was normalized to 18S using the 2^-ΔΔCT^ method. The primers used for both preamplification and qPCR are listed in Table 1.

**Table 1.**
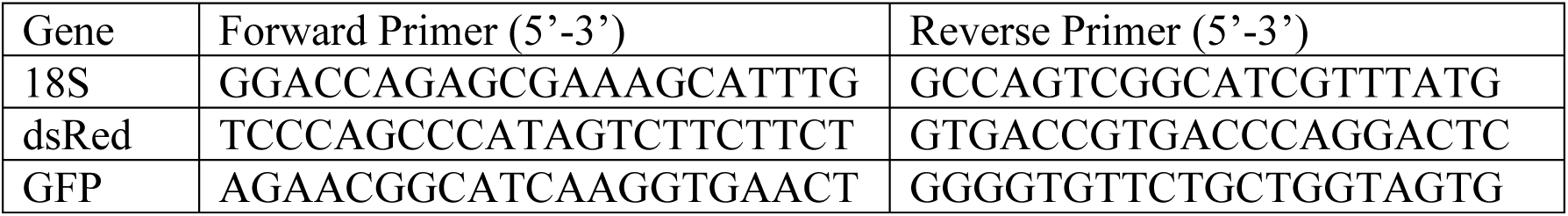
Primers used for cDNA preamplification and qPCR.

### Statistics

An unpaired t-test or one-way ANOVA with Bonferroni post hoc analysis were used for statistical evaluation. Data are expressed as the mean ± standard error (SE), and p < 0.05 was considered statistically significant. The number of replicates are as follows: RNA seq, 7-10; qPCR, 6; all other analyses, 3. Statistical analyses were performed using GraphPad Prism8 and R. Data values and p-values are reported within the text.

---

**Supplementary Fig. 1:**
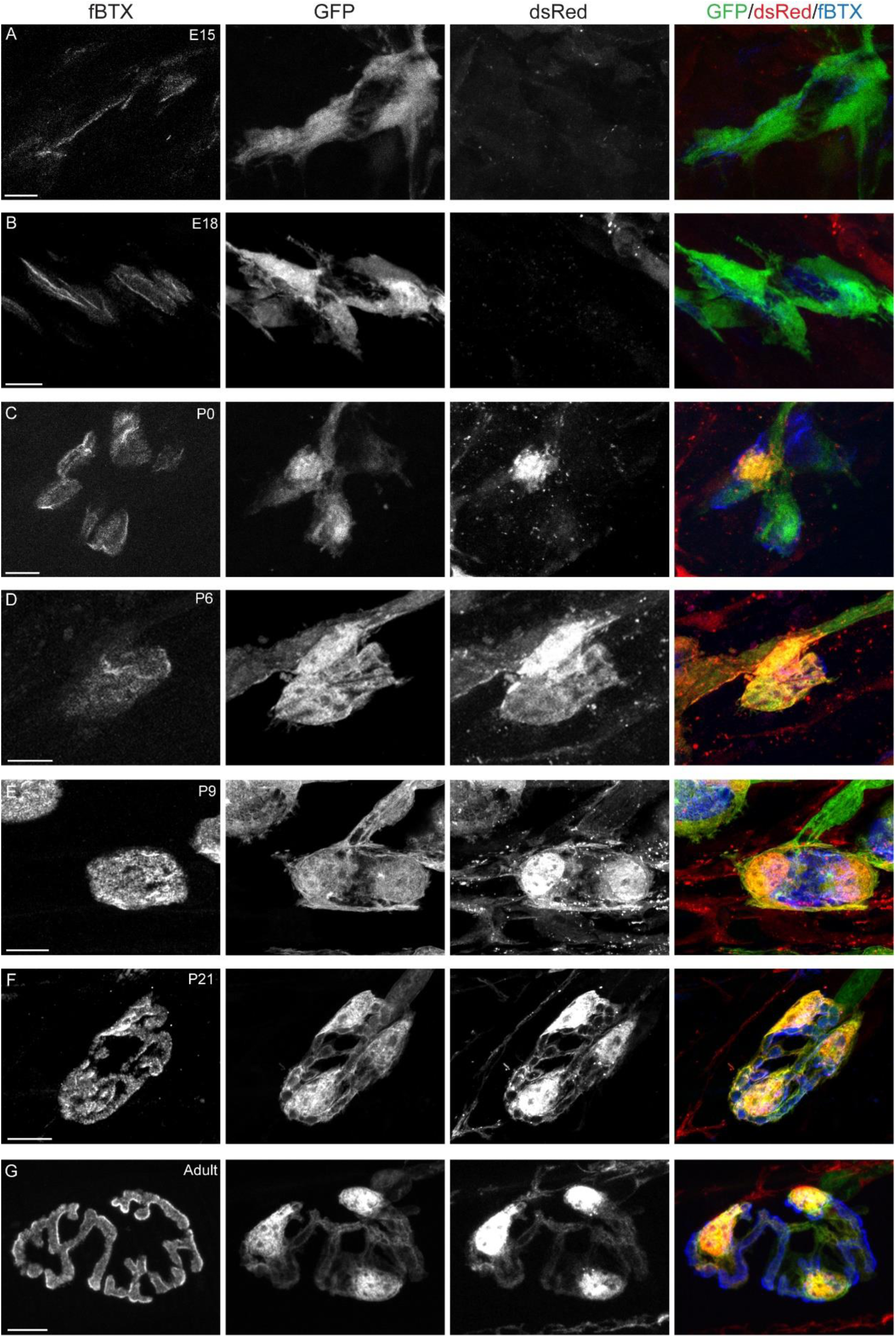
Color and grayscale images of PSCs at E15 (A), E18 (B), P0 (C), P6 (D), P9 (E), P21 (F), and adult (G). Scale bar = 10μm.

**Supplementary Fig. 2:**
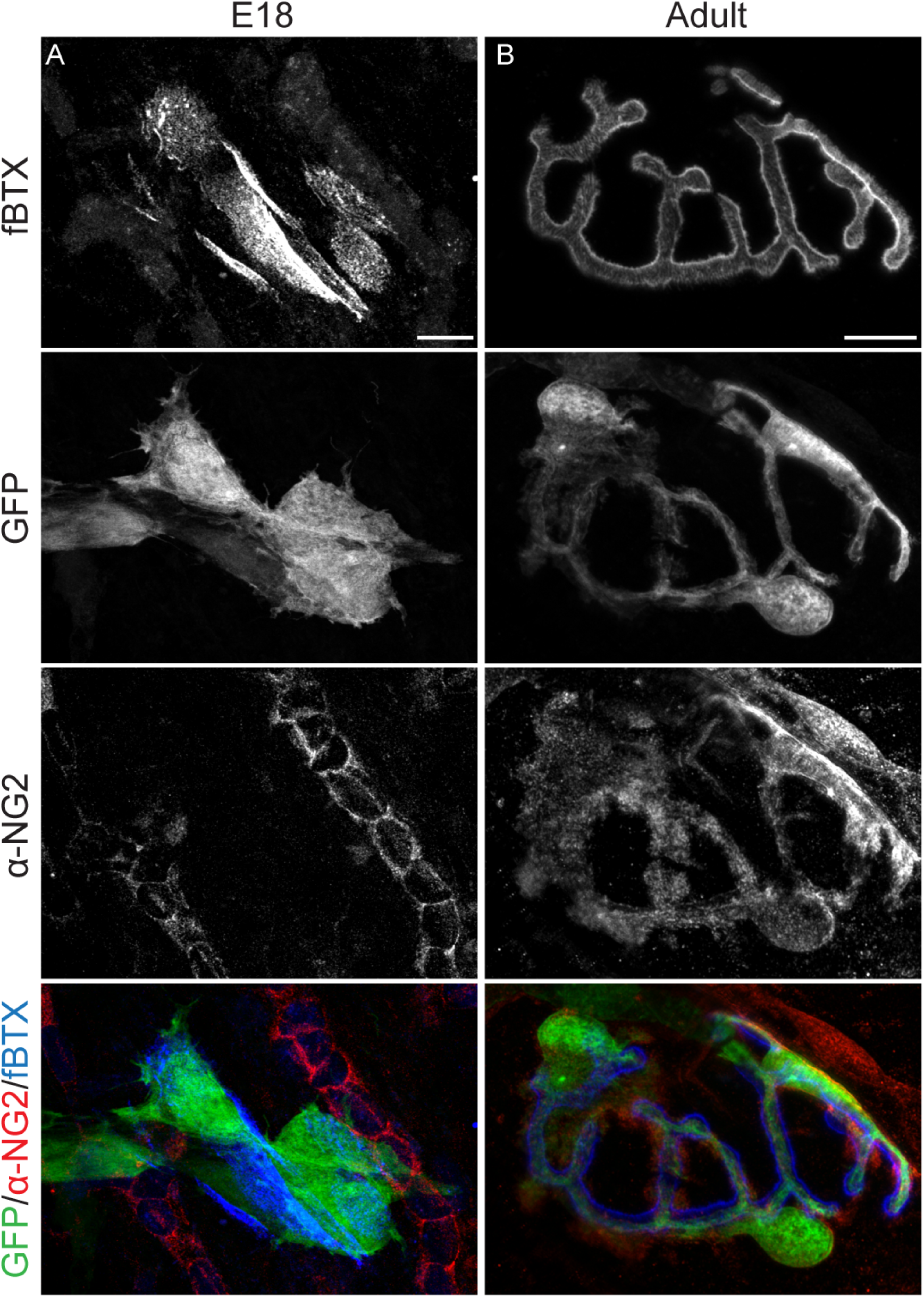
Cells at NMJs express NG2 in adults but not at embryonic timepoints. Immunohistochemical labeling of NG2 revealed that GFP+ cells at NMJs do not express NG2 in E18 mice (A). However, GFP+ cells at NMJs do express NG2 in adult mice (B). Scale bar = 10 μm.

**Supplementary Fig. 3:**
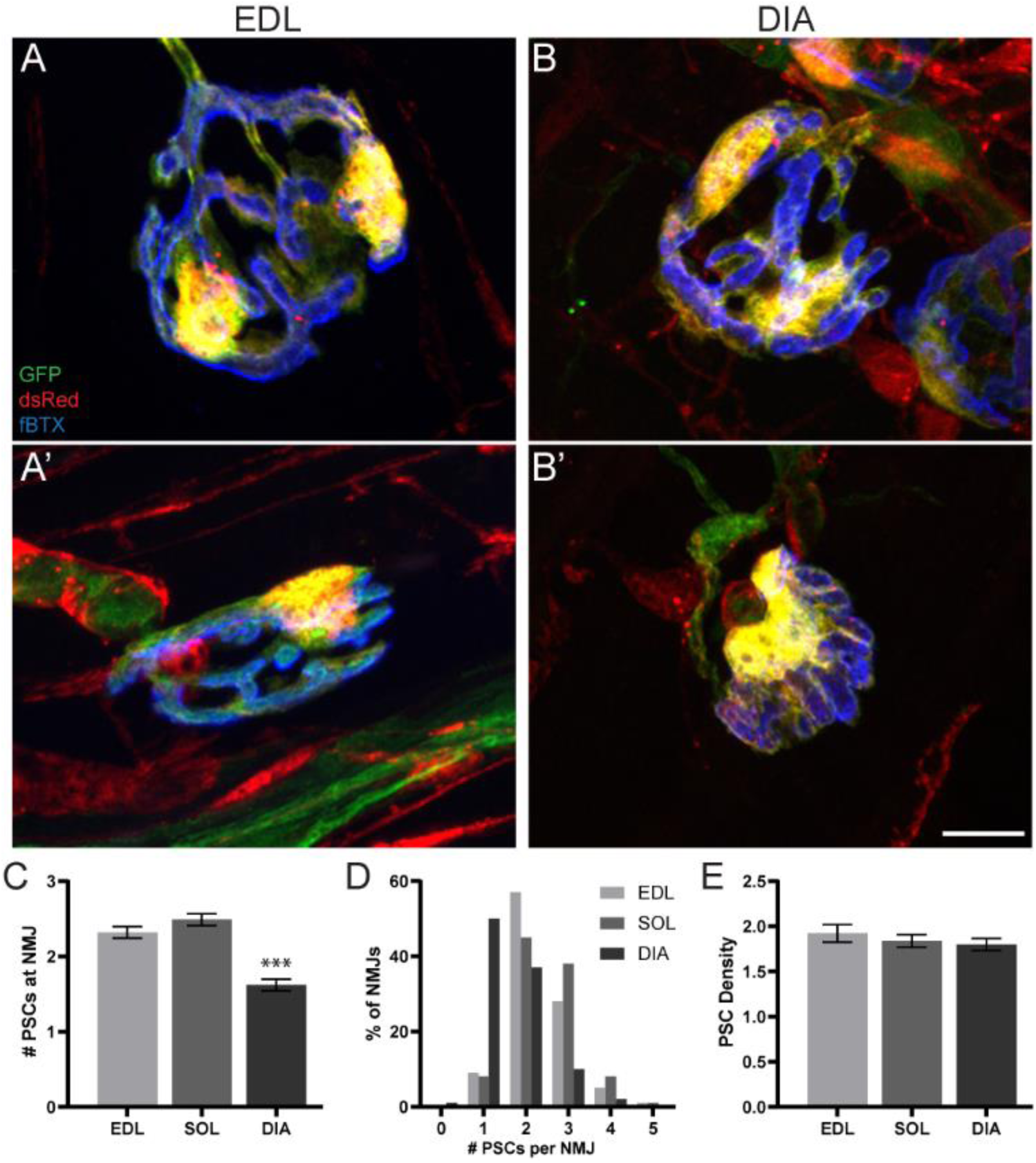
In the EDL, soleus, and diaphragm muscles of adult animals, the number of PSCs at NMJs varies (A-D). In each muscle, the number of PSCs per NMJ ranges from 0 to 5 PSCs per NMJ (D). When standardizing for NMJ size, the density of PSCs at NMJs is unchanged between muscles (E).

**Supplementary Fig. 4:**
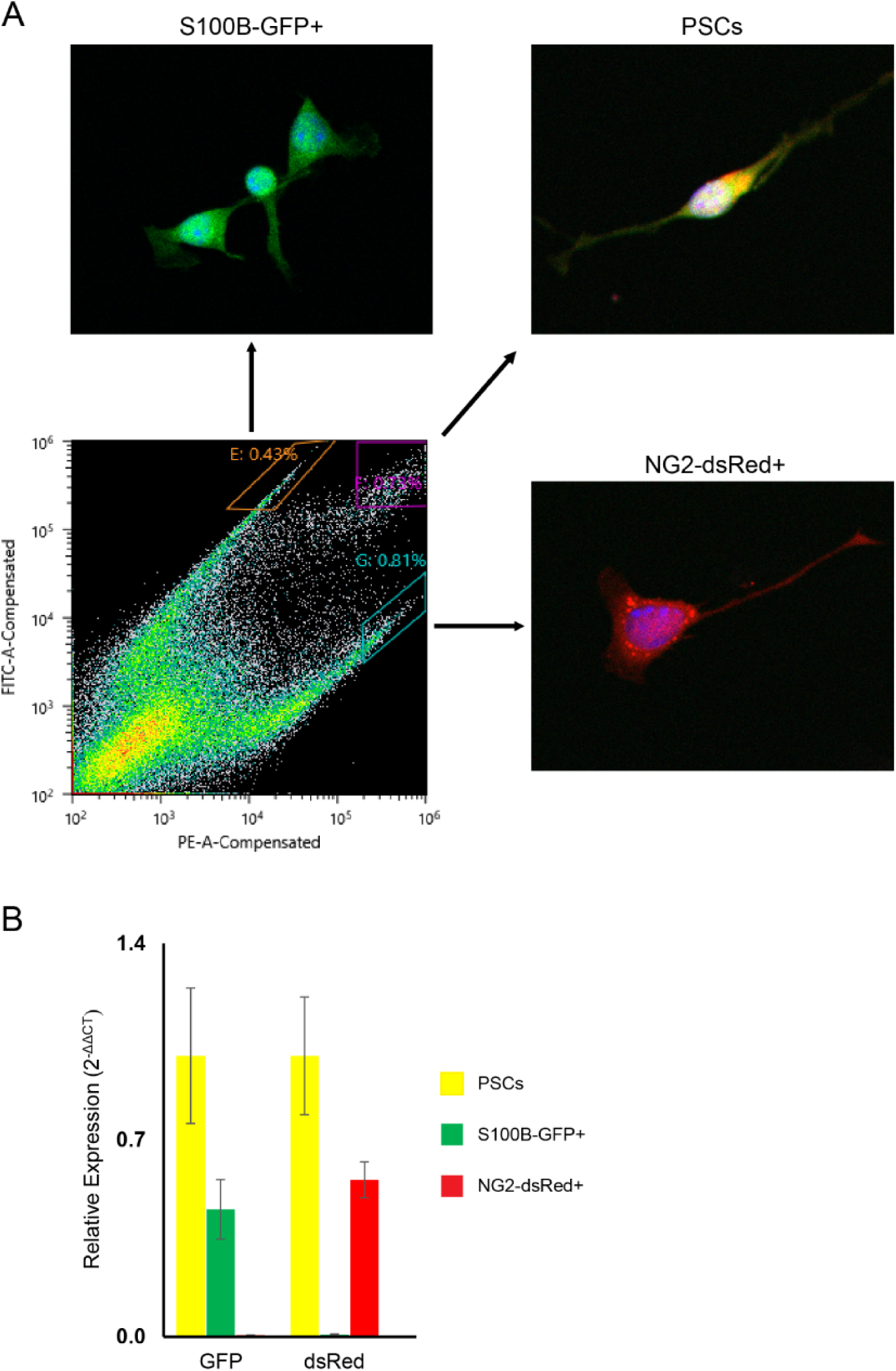
Representative images of fluorescence intensity gates and cells following FACS isolation of PSCs, S100B-GFP+, and NG2-dsRed+ cells from dissociated skeletal muscle tissue taken from S100B-GFP;NG2-dsRed mice (A). Confirmation of cell-specific dsRed and GFP expression with qPCR in PSCs, S100B-GFP+, and NG2-dsRed+ cells following FACS (B).

